# Beta-Cell Pyroptosis – A Burning Flame in Type 1 Diabetes?

**DOI:** 10.1101/2024.01.05.574294

**Authors:** Caroline Frørup, Cecilie Amalie Søndergaard Svane, Kristine Henriksen, Simranjeet Kaur, Joachim Størling

## Abstract

Immune-mediated destruction of the beta-cells in the pancreatic islets of Langerhans is the underlying cause of type 1 diabetes (T1D). Despite decades of research, the exact mechanisms involved at the beta-cell level during the development of disease remain poorly understood. This includes the mode(s) of beta-cell death and signaling events implicated in exacerbating local islet inflammation and immune cell infiltration, commonly known as insulitis. In disease models, beta-cell apoptosis seems to be the predominant cell death form which has led to the general assumption that beta cells mostly die by apoptosis in T1D. However, apoptosis is an anti-inflammatory programmed cell death mechanism, and this dogma is therefore challenged by the pathogenetic nature of T1D, as a progressive increase in islet inflammation is seen. This infers that other modes of beta-cell death that inherently increase insulitis may predominate. One such mechanism could be the newly characterized form of programmed cell death; pyroptosis (from the Greek “fire–falling”). Pyroptosis is characterized by gasdermin-mediated cell lysis with a bursting release of pro-inflammatory factors. Beta-cell death by pyroptosis in T1D may therefore offer a plausible explanation for the exacerbated paracrine islet inflammation that spreads during the progression of insulitis. Here, we briefly debate the evidence supporting beta-cell pyroptosis in T1D as a central mechanism of islet inflammation and beta-cell demise. The paper intends to challenge the current understanding of beta-cell destruction to move the field forward. Importantly, we present experimental data from human islets and EndoC-βH5 cells that directly support beta-cell pyroptosis as a rational death mechanism in T1D. We suggest a model of beta-cell demise in T1D in which pyroptosis plays a prominent role in concert with other cell death mechanisms. As the role of pyroptosis in disease is still in its infancy, we hope also to inspire researchers working in other disease fields.

## Introduction

Type 1 diabetes (T1D) is caused by autoimmune and inflammatory processes in the pancreas leading to destruction of the insulin-producing beta cells in the islets of Langerhans [1–3]. Its etiology is based on a combination of genetic predisposition and environmental triggering events such as nutritional components, bacterial and particularly viral infections have been suggested as possible initiators of the immunogenic reaction against the beta cells [4, 5]. Immune-cell infiltration of the islets and the release of pro-inflammatory factors, including cytokines and chemokines are hallmarks of T1D creating an inflammatory intra-islet milieu known as insulitis which may transpire for several years before clinical manifestation [6–12]. Interestingly, and for hitherto unknown reasons, insulitis tends to manifest as peri-insulitis, i.e., at one pole of the islet [13]. The discovery that residual beta cells persist years or even decades after disease onset in many patients [14–16] lends hope that new therapeutic approaches aiming at halting the beta-cell-destructive intra-islet processes will change diabetes trajectories.

One hindrance in arresting beta-cell death in T1D is that the nature by which the beta cells are lost i.e., the type(s) of death they undergo *in situ* remains largely undeciphered. The presumed dogma is that beta-cell death primarily happens by *apoptosis* [17, 18]. This anti-inflammatory form of cell death resolves in membrane blebbing and packaging of cellular contents in apoptotic bodies with ‘eat me’ flags for phagocytosis leaving the ‘crime scene’ in a non-inflamed and non-immunogenic state thereby preventing escalation and further inflammatory reactions [19, 20]. On this premise, beta-cell destruction in T1D should be quiescent, and non-immunogenic, thereby diminishing insulitis. However, during disease development, spatio-temporal peri-insulitis that gradually spreads within and between the islets, like a wildfire, is believed to take place [1, 13]. In accordance with this, studies have shown that blocking of apoptosis alone is inadequate in hindering beta-cell death in *in vitro* models [21–23]. Therefore beta-cell loss in diabetes cannot be explained by the current assumptions of the main beta-cell death mechanisms but leaves the mode(s) of human beta cell in T1D open for debate.

It has become clear that beta cells under immune attack play active roles in their own demise and are therefore not passive bystanders in the immune-mediated killing [24–26]. This is for instance exemplified by the active synthesis and release of multiple cytokines and chemokines from the beta cells upon immune attack causing detrimental auto- and paracrine effects, increasing immune cell co-activation, and chemoattractive immune effects [27].

Several regulated or programmed cell death mechanisms exist, and very likely beta cells die by several types of death mechanisms during T1D [17, 28, 29]. Mukherjee et al. recently thoroughly discussed the evidence for apoptosis, necrosis, and necroptosis in beta-cell loss in diabetes [17]. However, other forms of tightly controlled cell death exist, some of which remain entirely unresolved in diabetes and could contribute to beta-cell loss. In this article, we turn our attention to the plausible involvement of another more recently described programmed cell death form, i.e., *pyroptosis*. We summarize the current direct and circumstantial evidence advocating for pyroptosis as a possible beta-cell death form in human T1D and provide novel experimental data supporting this.

### Pyroptosis

Pyroptosis originates from the Greek *pyro* (fire) and *ptosis* (falling) and is a newly described form of programmed cell death, that is particularly pro-inflammatory by nature, resulting in the release of pro-inflammatory factors with a lytic endpoint [30, 31]. Physiologically, pyroptosis is a key cellular inflammatory response against extracellular pathogens like microbial and viral infections, ligands, or cellular perturbations [32]. However, pyroptosis is also known to play a critical role in various inflammatory diseases [33, 34]. Pyroptosis belongs to the innate immune response and can be classified as a cell death form orchestrated through canonical and/or non-canonical signaling pathways involving members of the gasdermin (GSDM) family. Canonical pyroptosis can be initiated by the binding of damage-associated molecular patterns (DAMPs), pathogen-associated molecular patterns (PAMPs) or homeostasis-altering molecular processes (HAMPs) to cell surface pattern recognition receptors (PRRs) of target cells. Stimulation of PRRs activates the inflammasome which consists of the nucleotide-binding domain, leucine-rich–containing family, pyrin domain–containing-1 and −3 (NLRP1/3) inflammasome, a sensor PRR, adaptor apoptosis-associated speck-like protein containing CARD (ASC), and pro-caspase-1. Once activated, caspase-1 processes the pro-inflammatory cytokines interleukin (IL)-1β and IL-18 into their mature forms and cleave GSDMD to its active form [35]. GSDMD then multimerizes and forms pores in the cell membrane that release mature IL-1β and IL-18, facilitate water influx, and hence, cell swelling ultimately resulting in lytic cell death. The release of inflammatory factors from cells undergoing pyroptosis is not restricted to IL-1β and IL-18, as other cytokines as well as chemokines are also released, including tumor necrosis factor-α (TNFα) and monocyte chemoattractant protein-1 (MCP-1) [36]. At the point of cell lysis, all cellular content including additional inflammatory factors is released. Non-canonical pyroptosis can be induced by e.g., bacterial lipopolysaccharide (LPS) through interferon-induced guanylate-binding proteins (GBPs) or directly by activating caspase-4/5/11 and subsequently GSDMD [35, 37, 38]. Other inducers of pyroptosis are granzyme A and B, secreted from cytotoxic lymphocytes, that cleave GSDMB [39] and GSDME, respectively, to activate pyroptosis [40]. Further, TNFα-stimulated caspase 8 activation can under certain conditions lead to cleavage of GSDMC thereby switching apoptosis to pyroptosis [41]. Likewise, caspase 3, the classical executioner caspase in apoptosis signaling, can also cleave and, hence, activate GSDME to induce pyroptosis suggesting a direct link between apoptosis and pyroptosis [42]. The GSDM-mediated release of cytokines and the eventual lytic release of a multitude of cellular contents including proteases, cytokines, and chemokines exacerbate local tissue inflammation and cause paracrine spreading of pyroptosis [35, 37, 43].

### New perspectives on pyroptosis in type 1 diabetes

Emerging evidence indicates that pyroptosis of beta cells may be involved in the pathogenesis of diabetes. Thus, in a recent study, we identified *GSDMB* as one of only few common candidate risk genes for T1D and type 2 diabetes (T2D) that likely operate at the islet level [44]. This infers that part of the genetic susceptibility underlying both forms of diabetes could be due to altered GSDMB-dependent pyroptosis signaling. Interestingly, we found islet expression quantitative trait locus (eQTL) signals in linkage disequilibrium (LD) with variants associated with T1D and T2D for *GSDMB* [44], suggesting that genetic variants affecting the expression level of *GSDMB* in islets might influence the beta cells’ sensitivity to pyroptotic cell death.

Presently, only a few studies have thus far investigated pyroptosis in beta cells and islets [45, 46]. It was recently reported that elevated blood glucose levels induce beta-cell pyroptosis in diabetic mice, while reduced pyroptosis coincides with lower blood glucose levels [46, 47]. Liu et al. showed that pyroptosis signaling events were increased in response to elevated glucose in the rat beta-cell line INS-1 and that GSDMD cleavage was increased in streptozotocin-treated diabetic mice [45]. This was recently confirmed in a study by Zhou et al. who found that high glucose alone or combined with high insulin increased pyroptosis in a time-dependent manner in INS-1 cells. They showed that salidroside, a plant glucoside, and a small molecule inhibitor of NLRP3 were able to reverse the increase in pyroptosis markers in the pancreas of streptozotocin-treated diabetic mice [46]. Opposed to these findings, Wali et al. reported that genetic ablation of NLRP3 or caspase-1 did not protect against oxidative, endoplasmic reticulum stress, or glucotoxicity in isolated mouse islets [48]. In INS-1 cells, Ghiasi et al. reported that both *Nlrp1* and *Nlrp3* were modulated in response to cytokine exposure and showed that a broad inflammasome inhibitor protected against IL1β-induced toxicity, which was not observed upon NLRP3-selective inhibition [49].

Two studies have shed light on pyroptosis in isolated human islets and pluripotent stem cell-derived beta-like cells. In one study, the authors reported that *GSDMB*, *GSDMD*, *GSDME*, caspase 1 (*CASP1*), and 4 (*CASP4*) were significantly upregulated in human islets and human beta-like cells after treatment with reactive chimeric antigen receptor (CAR) T-cell conditioned medium [50] indicating a putative role of beta-cell pyroptosis in immune-mediated beta-cell killing in T1D. Another study reported the expression of several inflammasome-related genes (IRGs) in human islets among which mitogen-activated protein kinase 8 interacting protein-1 (MAPK8IP1) was suggested to regulate several IRGs, including GSDMD based on siRNA-silencing experiments in INS-1 cells [51]. Lebreton et al. reported that exposure to LPS and successively to ATP upregulated *NLRP3* in human islets correlating with the processing of pro-IL-1β to mature IL-1β [52]. These studies collectively support that key markers of pyroptosis and upstream signaling components are induced in beta cells in experimental models, inferring that beta-cell pyroptosis could be an important cell death mechanism in diabetes. However, human beta-cell data are scarce and the full implication of pyroptosis therefore await further clarification.

To provide further information about alterations of pyroptosis-related genes in human beta cells in the context of diabetes, we used available human islet single cell RNA sequencing data from the human pancreas analysis program (HPAP, http://www.tools.cmdga.org:3838/isletHPAP-expression/ [53] to extract beta-cell gene expression of several pyroptosis-related genes. The heatmap in Figure 1A shows that the expression of several pyroptosis-related genes including *GSDMD*, *CASP1*, and *NLRP1/3* is increased in beta cells from donors with T1D. Interestingly, *GSDMB* and *GSDME* are more highly expressed in beta cells from autoantibody-positive donors compared to beta cells from T1D donors. Also, beta cells from T2D donors show higher expression of *GSDMB* and *GSDMC* compared to beta cells from T1D donors. This may reflect different roles of gasdermins in beta-cell inflammation during development of T1D and during sustained inflammation in type 2 diabetes versus established T1D where most of the beta cells have perished. We also examined if pro-inflammatory cytokines, known as key drivers of beta-cell damage and death, upregulate pyroptosis genes in human islets and human EndoC-βH5 beta cells [54]. The heatmaps in Figure 1B and C generated from in-house RNA sequencing data show that cytokines increase the expression of several pyroptosis genes in human islets and EndoC-βH5 cells. Real-time qPCR of selected key genes validated the RNA sequencing data (Figure 1D and E). Importantly, we confirmed upregulation of GSDMD at the protein level in response to cytokine treatment in EndoC-βH5 cells and human islets by Western blotting (Figure 1F and G). Together, these observations add evidence for a role of pyroptosis in human beta cells in T1D *in situ* and *ex vivo*.

**Figure 1:**
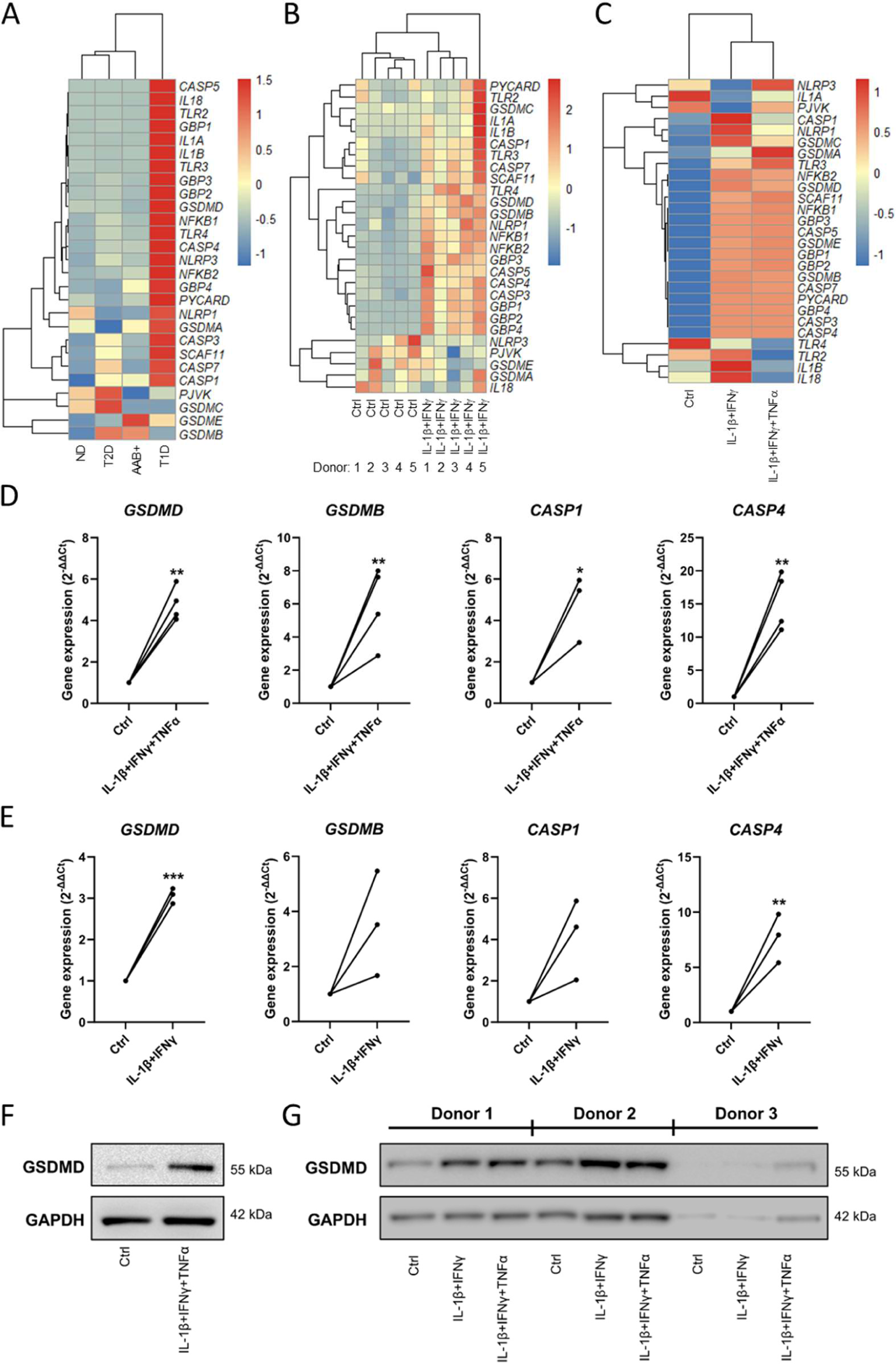
Increased expression of pyroptosis genes in beta cells from donors with T1D, and upregulation of pyroptosis genes by pro-inflammatory cytokines. **A**) Heatmap of expression of pyroptosis-related genes in beta cells from donors with T1D, T2D, autoantibody positive donors (AAB+), or non-diabetic (ND) donors. Data grouped by diabetes status were extracted from the HPAP dataset (http://www.tools.cmdga.org:3838/isletHPAP-expression/). The normalized TPM (transcripts per million) expression values plotted as scaled gene expression (z-scores) with gene and sample clustering. **B**) and **C**) Heatmaps of expression of pyroptosis-related genes in human islets and EndoC-βH5 beta cells, respectively. Scaled TPM values from 5 individual non-diabetic human islet donors are shown. For EndoC-βH5, scaled mean TPM values of n=4 are plotted. Untreated control (Ctrl) or exposed to cytokines (human islets: 50 U/ml interleukin-1β (IL-1β) + 1,000 U/ml interferon-γ (IFNγ) for 24 hours; EndoC-βH5 cells: 50 U/ml IL-1β + 1,000 U/ml IFNγ ± 1,000 U/ml tumor necrosis factor α (TNFα) for 48 hours). All heatmaps created using the pheatmap package in RStudio. **D**) and **E**) Expression of selected pyroptosis genes determined by real-time PCR using TaqMan assays in EndoC-βH5 cells (panel D, n=4) and human islets (panel E, n=3). Gene expression was normalized to *GAPDH* expression. **F**) and **G**) Western blot analysis of extracts from EndoC-βH5 cells (n=4) and human islets from three non-diabetic donors (n=3), respectively, using an antibody against GSDMD (cat no. #39754, Cell Signaling Technology). Islets and cells were exposed to cytokines for 24 and 96 hours, respectively. GAPDH was used as loading control. Representative blots are shown. *: p<0.05, **: p<0.01, ***p<0.001, paired *t*-test.

### Suggested model of beta-cell pyroptosis in type 1 diabetes

The data presented above together with the reviewed literature favor that beta-cell pyroptosis could play an important role in the active signaling of programmed beta-cell death and may thus be a hitherto underrecognized form of beta-cell death. As opposed to apoptotic cell death, pyroptosis can explain key hallmarks of the disease including secretion of beta-cell pro-inflammatory cytokines and chemokines, exacerbation of insulitis, and the spreading of beta-cell inflammation like a fire across the islet [1]. We propose a model (Figure 2) where pyroptosis contributes to and accelerates beta-cell destruction by being a key driver of beta-cell inflammation. The model includes triggering events that have previously been coupled to both T1D and pyroptosis signaling e.g., viral and bacterial infection in seemingly arbitrary beta cells and islets. This is followed by immune-cell infiltration, beta-cell peptide presentation on MHC molecules, T-lymphocyte activation, and subsequent granzyme/perforin release. In the beta cell, synthesis, processing, and release of cytokines and chemokines firstly via gasdermin pores, secondly after cell lysis fuels aggravation and paracrine spreading of inflammation.

**Figure 2:**
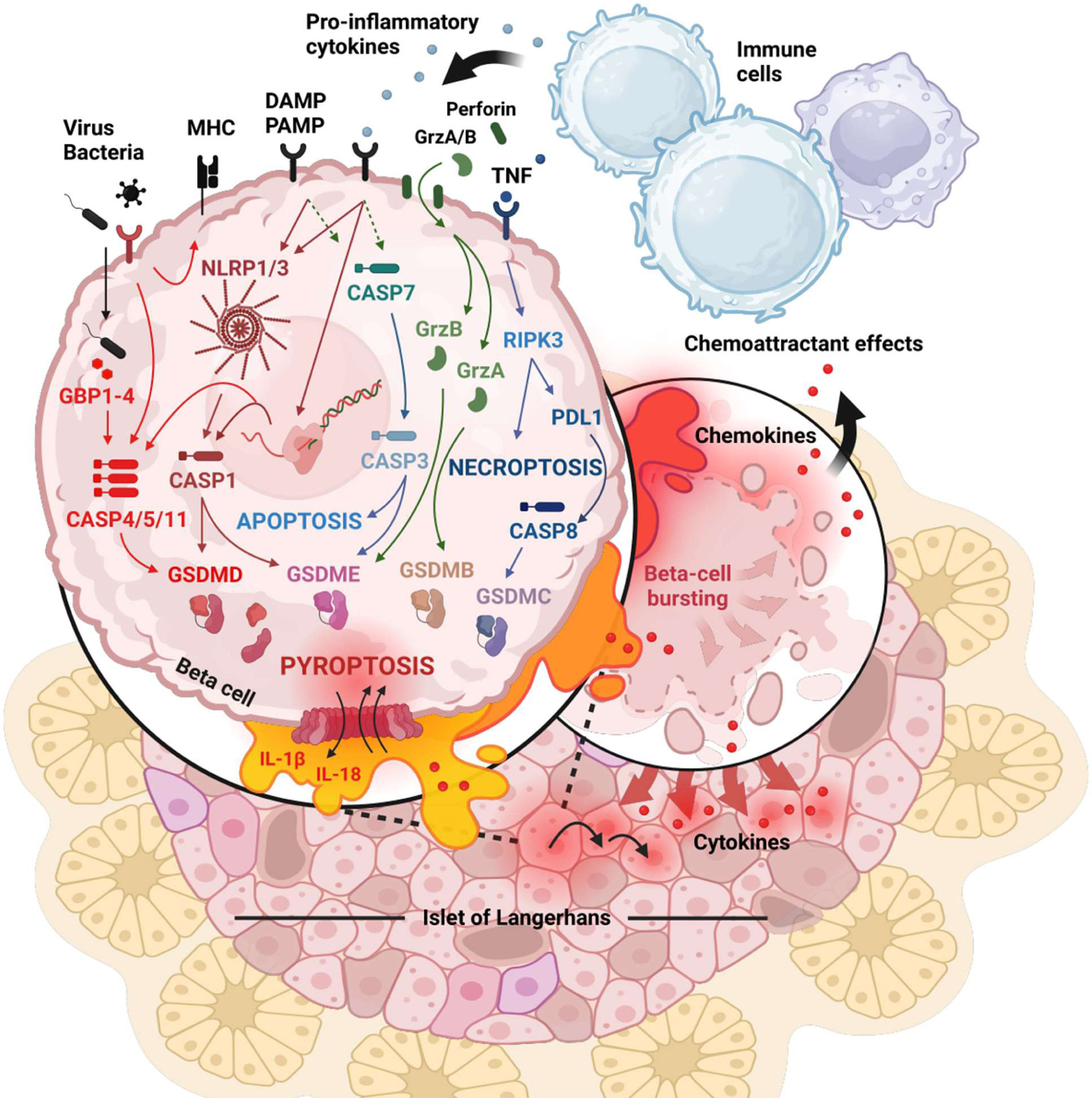
Hypothetical model of beta-cell pyroptosis in T1D. Simplified illustration suggestive of the potential for gasdermin-dependent pyroptosis along with other cell death types in the beta cells. Beta-cell pyroptosis is triggered by receptor activation and/or intracellular mechanisms culminating in caspase-mediated gasdermin cleavage and pore formation leading to release of pro-inflammatory factors (cytokines and chemokines) and ultimately cell lysis. This causes paracrine spreading and amplification of destructive islet inflammation processes including increased immune cell recruitment and activity. Figure was created in BioRender.com. CASP, caspase; DAMP/PAMP, damage/pathogen-associated molecular pattern; GSDM, gasdermin; GBP1, interferon-induced guanylate-binding protein 1; Grz, granzyme; IL, interleukin; MHC, major histocompatibility complex; NLRP, nucleotide-binding domain, leucine-rich–containing family, pyrin domain–containing-3; PAMPs, pathogen associated molecular patterns; PDL1, programmed death ligand 1; RIPK3; receptor-interacting serine-threonine kinase 3, TNF, tumor necrosis factor.

The model offers a plausible explanation for some of the yet unexplained phenomena of beta-cell death and T1D etiology compared to other cell death mechanisms including the spatio-temporal peri-insulitis occurring during disease development [1]. We suggest an interplay between several cell death mechanisms, including caspase-dependent apoptosis and pyroptosis, as well as caspase-independent necroptosis, exemplifying the complex nature of beta-cell demise in T1D.

### Present and Future Perspectives

Decades of research has advanced our insight into the immune-mediated killing of the beta cells characteristic of T1D, however, the exact intrinsic death mechanisms by which human beta cells perish in T1D remain obscure. The evidence and data discussed here support a relationship between pyroptosis signaling pathways and human beta-cell death, which could constitute a ‘missing link’ in the hitherto devised roadmap to beta-cell loss in T1D. Research in this area is in demand and may lay the foundation for a better understanding of disease etiology. In addition to being an important mediator of pyroptotic cell death, GSDMD also holds potential as a circulating biomarker, as serum GSDMD levels are elevated in inflammatory diseases [55–57]. Whether GSDMD or other gasdermins can serve as circulating biomarkers of ongoing beta-cell pyroptosis should be investigated. It remains to be proven that pyroptosis contributes to immune-mediated beta-cell destruction in T1D and models hereof, but if so, therapeutics targeting pyroptotic components could be used to prevent or treat T1D. Efforts set out to test this possibility in human cellular and preclinical models should be of high priority.

Changing the trajectory of T1D by halting beta-cell loss, proves difficult if we do not challenge current knowledge and see in new directions for opportunities. In this article, we have proposed that beta-cell pyroptosis should be considered a new important beta-cell death form in T1D offering a rational explanation of ‘loose ends’ in our current disease understanding. We advocate for further studies and avenues investigating pyroptosis as a key inflammatory beta-cell death mechanism in human T1D which can pave the way for new treatment strategies.

## Abbreviations

eQTL: expression quantitative trait locus
CAR: chimeric antigen receptor
CASP: caspase
DAMPs: damage-associated molecular patterns
GSDM: gasdermin
GBP1: interferon-induced guanylate-binding protein 1
HAMPs: homeostasis-altering molecular processes
IL-1β: interleukin-1β
IL-18: interleukin 18
IFNγ: interferon-γ
LD: linkage disequilibrium
NLRP: nucleotide-binding domain leucine-rich–containing family pyrin domain–containing
NOD: non-obese diabetic
PAMPs: pathogen-associated molecular patterns
PDL1: programmed death ligand 1
PPRs: pattern recognition receptors
RIPK3: receptor-interacting serine-threonine kinase 3
STZ: streptozotocin
TNFα: tumor necrosis factor alpha.

## Acknowledgments

We thank Caroline Lindgreen for excellent laboratory assistance.

## Author Contributions

C.F. and J.S. conceptualized and designed the study. C.F. made the initial manuscript draft. All authors contributed to the acquisition of data, critical scientific input, and data interpretation. The final version of the manuscript was approved by all authors.

## Ethics Approval and Consent to Participate

Isolated human islets from 5 non-diabetic donors were purchased from Prodo Laboratories Inc., Aliso Veijo, CA, USA. Donor information is presented in Supplementary Table 1.

## Data Availability Statement

All data are publicly available (http://www.tools.cmdga.org:3838/isletHPAP-expression/, Gene Expression Omnibus (GEO) repository accession number GSE218735) or available upon request.

---

**Supplementary Table 1.** Checklist for reporting human islet preparations used in research. Adapted from Hart NJ, Powers AC (2018) Progress, challenges, and suggestions for using human islets to understand islet biology and human diabetes. Diabetologia https://doi.org/10.1007/s00125-018-4772-2. <u>Human islets were purchased from Prodo Laboratories Inc, Aliso Veijo, CA, USA.</u>

| Islet preparation | 1 | 2 | 3 | 4 | 5 |
| --- | --- | --- | --- | --- | --- |
| Unique identifier | HP-19003-01 | HP-19073-01 | HP-19136-01 | HP-19150-01 | HP-19163-01 |
| Donor age (years) | 63 | 57 | 34 | 38 | 24 |
| Donor sex (M/F) | F | M | M | F | M |
| Donor BMI (kg/m<sup>2</sup>) | 27.0 | 35.9 | 28.0 | 24.1 | 24.5 |
| Donor HbA<sub>1c</sub> or other measure of blood glucose control | 5.8 | 5.4 | 5.2 | 5.8 | 5.6 |
| Donor history of diabetes?<br>Please select yes/no from drop down list | No | No | No | No | No |
| Donor cause of death | Stroke | Stroke | Anoxic event | Anoxic event | Cerebrovascular accident |
| Estimated purity (%) | 85 | 90 | 90 | 85-90 | 95 |
| Estimated viability (%) | 95 | 95 | 95 | 95 | 95 |

